# The interfascicular matrix of energy storing tendons houses heterogenous cell populations disproportionately affected by ageing

**DOI:** 10.1101/2023.01.04.522701

**Authors:** Danae E. Zamboulis, Neil Marr, Luca Lenzi, Helen L. Birch, Hazel R. C. Screen, Peter D. Clegg, Chavaunne T. Thorpe

## Abstract

Energy storing tendons such as the human Achilles and equine superficial digital flexor tendon (SDFT) are prone to injury, with incidence increasing with ageing. The interfascicular matrix (IFM), which binds tendon fascicles, plays a key role in energy storing tendon mechanics, and ageing alterations to the IFM negatively impact tendon function. While the mechanical role of the IFM in tendon function is well-established, the biological role of IFM-resident cell populations remains to be elucidated. Therefore, the aim of this study was to identify IFM-resident cell populations and establish how these populations are affected by ageing. Cells from young and old SDFTs were subjected to single cell RNA-sequencing, and immunolabelling for markers of each resulting population used to localise cell clusters. Eleven cell clusters were identified, including tenocytes, endothelial cells, mural cells and immune cells. One tenocyte cluster localised to the fascicular matrix, whereas nine clusters localised to the IFM. Interfascicular tenocytes and mural cells were preferentially affected by ageing, with differential expression of genes related to senescence, dysregulated proteostasis and inflammation. This is the first study to uncover the importance of the IFM niche for a diverse range of cell populations, and to identify age-related alterations specific to IFM-localised cells.

## Introduction

Tendons transmit muscle-generated force to the skeleton, and specific tendons also play a key role in locomotory efficiency by storing and returning energy with each stride. These energy storing tendons are prone to age-related degeneration and subsequent injury (Knobloch et al., 2008, Clayton and Court-Brown, 2008, Perkins et al., 2005), however the causes of this degeneration remain to be established. The human Achilles tendon and equine superficial digital flexor tendon (SDFT) are the predominant energy storing tendons in the human and horse respectively (Biewener, 1998, Lichtwark and Wilson, 2007). Indeed, the equine SDFT is a relevant and well-established model in which to study tendon ageing, due to similarities in structure and function with the human Achilles, as well as shared epidemiology and aetiology of disease, and poor healing after injury (Patterson-Kane and Rich, 2014, Innes and Clegg, 2010).

Tendons are rich in extracellular matrix proteins, predominantly collagen type I, which is arranged in a hierarchical manner and interspersed with non-collagenous proteins (Kastelic et al., 1978). At the largest level of the hierarchy, tendon fascicles are bound by the interfascicular matrix (IFM), a looser connective tissue matrix rich in proteoglycans, minor collagens and elastin (Thorpe et al., 2016b, Thorpe et al., 2016a, Godinho et al., 2021, Godinho et al., 2017). Cells, collectively referred to as tenocytes, are found in both the fascicles and IFM regions, with greater cell density within the IFM (Thorpe et al., 2016b). There is also evidence of other cell populations present within tendon, with single-cell RNA sequencing (scRNAseq) of murine and human tendon revealing several tenocyte populations, as well as endothelial, mural and immune cell populations (Kendal et al., 2020, De Micheli et al., 2020). However, the tendons used in these studies generally have limited energy storing capacity and a less prominent IFM, therefore it has not yet been established which cell populations localise to the IFM, nor how tendon cell populations may be differentially affected by ageing.

The IFM plays a key role in the function of energy storing tendons by allowing sliding and recoil between fascicles, providing the whole tendon with high strain capacity and fatigue resistance (Thorpe et al., 2012, Thorpe et al., 2015, Thorpe et al., 2016c). The IFM is disproportionately affected by ageing, with IFM stiffening, reduced fatigue resistance and impaired recoverability reported, all associated with an increased risk of injury in aged tendon (Thorpe et al., 2013, Thorpe et al., 2017). Therefore, characterising the role of IFM cell populations in maintaining tendon homeostasis, and establishing how IFM cell populations are impacted by ageing is crucial to elucidate the drivers of age-related alterations in tendon structure-function relationships and how these lead to increased risk of injury with ageing. Therefore, the aim of this study was to use single-cell RNA sequencing, combined with immunolabelling, to identify and localise IFM-resident cell populations in the equine SDFT and establish how these populations are affected by ageing.

## Results

### 11 cell clusters are present in equine tendon

To characterise young and aged tendon populations in an energy storing tendon, scRNA-seq was performed on SDFT cells isolated from 4 young and 4 old horses (Figure S1). Following quality control and filtering, data were derived from a total of 59273 cells which clustered into 11 distinct clusters (Figure 1A). Three tenocyte clusters were identified based on expression of *COL1A1, COL3A1, COMP, DCN, LUM* and *LOX*. The largest of these clusters was defined as FM tenocytes (FM), based on higher levels of expression of *COMP, LOX* and *THBS4* (Figure 1B), which are localised to the fascicular matrix (Södersten et al., 2013, Frolova et al., 2014). The second tenocyte cluster was defined as IFM tenocytes (IFM) due to their differential expression of *PRG4*, a well-established marker of IFM cells (Thorpe et al., 2016a, Kohrs et al., 2011, Sun et al., 2015), and *TNXB* (Figure 1B), localisation of which was confirmed using immunolabelling (Figure 1C). The third tenocyte cluster expressed markers found in both IFM and FM tenocyte clusters (*COMP, LOX, THBS4, TNXB, LUM, PRG4)* and is referred to as mixed tenocytes (MixT). Clustering of tenocytes in the “MixT” cluster was driven by significantly lower expression of ribosome biogenesis-specific genes (Figure S2A), which was more pronounced in aged tendons. Two mural cell clusters (MuC1&2) were identified which localised to the IFM region, based on differential expression of the vascular smooth muscle and pericyte markers *RGS5, MYH11* and *ACTA2* (Vanlandewijck et al., 2018, Muhl et al., 2020) (Figure 1B). Immunolabelling for these markers revealed the mural cells surrounded vessels in the IFM and were absent in the fascicles (Figure 1C). Similar to the MixT cluster, clustering of mural cells in the MuC2 cluster was driven by lower expression of ribosomal biogenesis genes (Figure S2B). Vascular endothelial cells, (EC_V) expressing *PECAM1, VWF* and *SELE*, and lymphatic endothelial cells (EC_L), expressing *PECAM1, MMRN1* and *CCL21* also localised to the IFM region (Figure 1B and C). Four immune cell clusters were identified, based on expression of *PTPRC* and *CD74*, which localised to the IFM (Figure 1B and C). These clusters were further identified as T cells (TC; *CCL5* and *CD3E*), macrophages (Mφ; *CSF1R* and *G0S2*), neutrophils (Neu; *MSR1* and *CSF1R*) and mast cells (MC; *KIT*) (Figure 1B).

**Figure 1.**
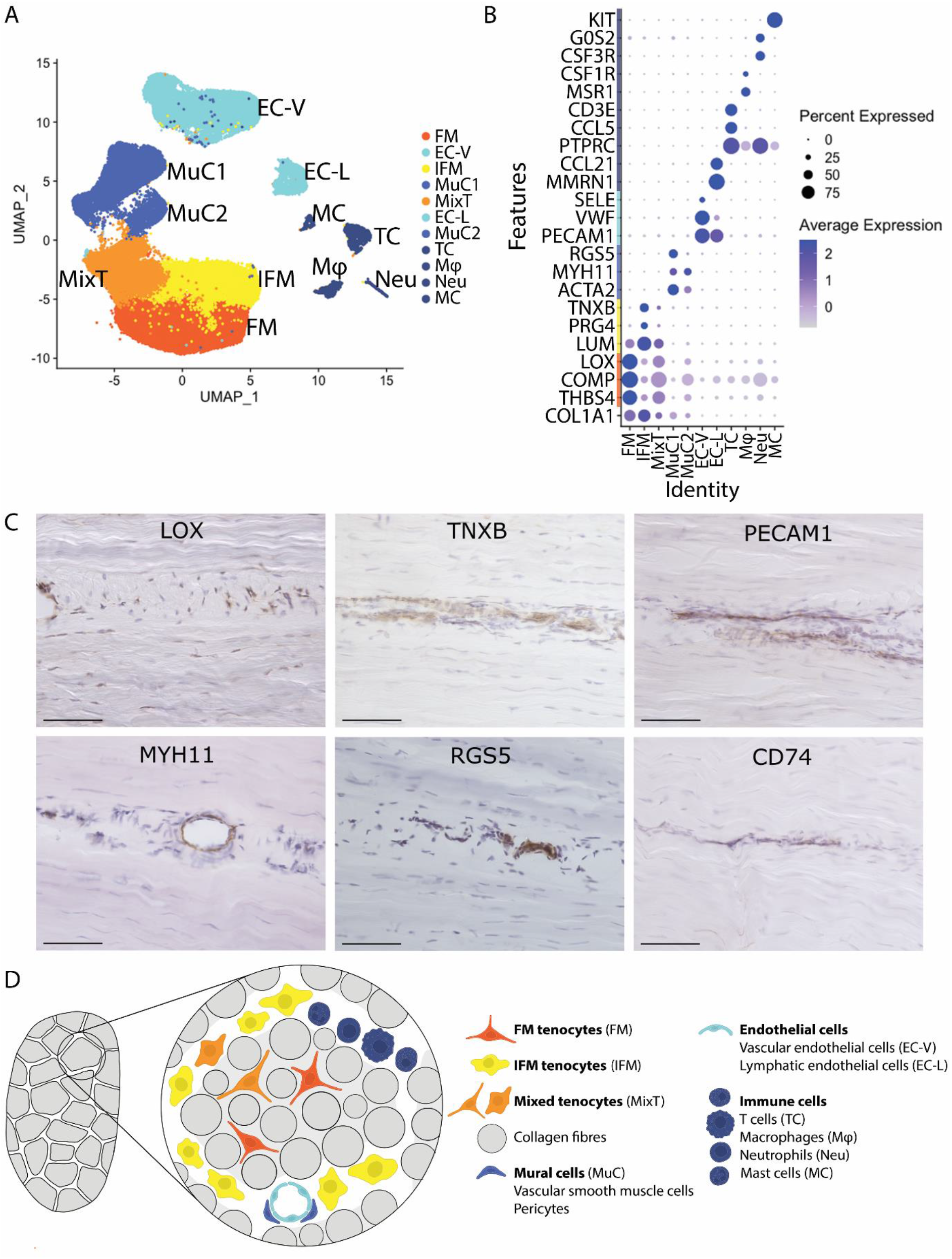
11 cell clusters are present in equine tendon. (A) Uniform Manifold Approximation and Projection (UMAP) dimensionality reduction demonstrates the presence of 11 clusters, based on differential gene expression, namely FM tenocytes “FM”, IFM tenocytes “IFM”, mixed tenocytes “MixT”, mural cells “MuC1” and “MuC2”, vascular endothelial cells “EC-V”, lymphatic endothelial cells “EC-L”, T cells “TC”, macrophages “Mφ”, neutrophils “Neu”, and mast cells “MC”. Each cluster consists of cells originating from young and old donors (n=8). (B) Dot plot showing genes used to differentiate and identify the clusters. Scale indicates average expression and ranges from grey = 0 to blue = 2, dot size indicates the percentage of cells expressing the gene. (C) Immunolabelling of longitudinal tendon sections reveal that all cells within fascicles, and some within the IFM compartment are positive for LOX, and cells within the IFM compartment show presence of TNXB, the endothelial cell marker PECAM1, the mural cell markers MYH11 and RGS5 and the immune cell marker CD74. (D) Schematic of tendon demonstrating location of the different cell populations identified.

### IFM tenocytes and mural cells are preferentially affected by ageing

To establish the effect of ageing on tendon cell populations, the size of each cluster and the number of differentially expressed (DE) genes in each cell cluster from young and old tendons was assessed. The number of cells in MuC2, as a proportion of total cell number, increased significantly with ageing, whereas the number of Mφ, as a proportion of total cell number, decreased with ageing. Many genes within each cluster were also DE with ageing (Figure 2A and B). Clusters IFM and MuC2 were disproportionately affected by ageing, with 466 and 902 DE genes respectively. The Aging Atlas (Consortium, 2020) was used to identify DE genes in each cluster associated with ageing-related dysfunction. Many of the DE ageing genes were associated with senescence and senescence-associated secretory phenotype, as well as loss of proteostasis (Figure 2C), particularly in IFM and MuC2 clusters (Figure 2D). Further, comparison of the top 25 markers of each cluster (most highly expressed cluster differentiating genes) between young and old samples revealed age-dependent loss of the top genes characterising each cluster for the IFM, MixT and MuC2 clusters (Figure 2E). Many genes associated with inflammation (GOTerm: inflammatory response, GO:0006954) (Smith and Eppig, 2009) were also DE with ageing, particularly in IFM and MuC2 clusters (Figure 2F). Ageing also had an effect on cell cycling for the IFM cluster only where there was a significantly larger percentage of cells classifying in S phase in old samples compared to young samples (Figure S3).

**Figure 2.**
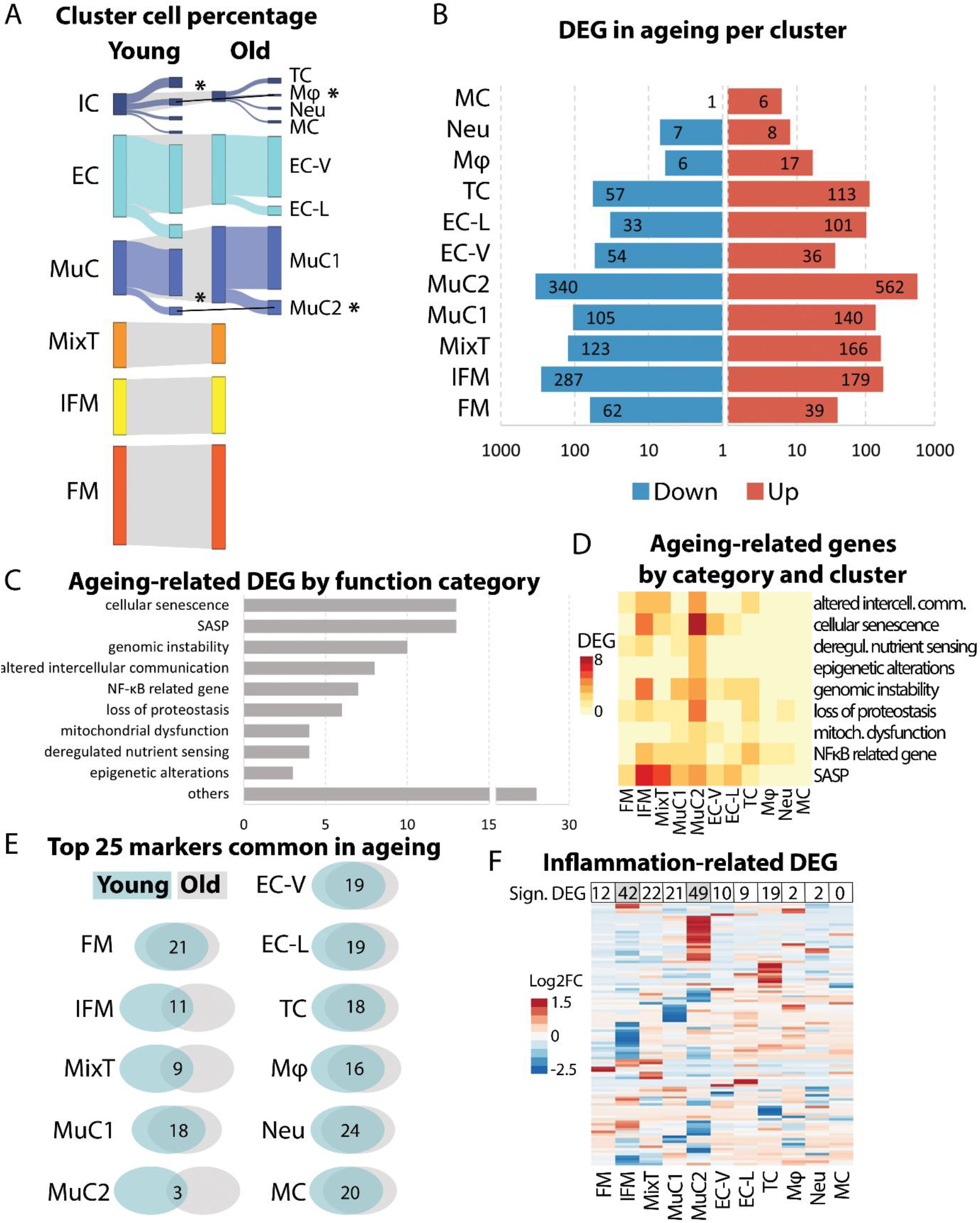
IFM tenocytes and mural cells are preferentially affected by ageing. (A) The percentage of cells in the majority of clusters was predominantly unaffected by ageing, except for an increase in the proportion of cells in mural cell cluster 2 (MuC2; p=0.049, t-test) and a decrease in the proportion of macrophages (Mφ; p=0.039, t-test). Significance is indicated by*. The number of differentially expressed genes (DEGs) with ageing varied between clusters, with the highest number of DEGs in MuC2 and IFM clusters. Data are plotted on a log_10_ scale. Ageing-related DEGs (Aging Atlas (Consortium, 2020)) between young and old tendon cells were primarily associated with senescence and the senescence-associated secretory pathway (SASP). (D) Heatmap demonstrating IFM tenocyte and MuC2 clusters had the greatest number of ageing related DEGs and in particular DEGs associated with senescence and SASP. Scale indicates number of genes and ranges from yellow = 0, to red = 10. (E) Venn diagrams showing the number of top 25 markers in each cluster that are maintained with ageing. An age-dependent loss of the top genes characterising each cluster is observed for the IFM tenocyte, MixT and MuC2 clusters. (F) Heatmap showing IFM tenocyte and MuC2 clusters had most DE inflammation-related genes (GOTerm: inflammatory response, GO:0006954) with ageing. Scale indicates log_2_FC and ranges from blue = –2.5, to white = 0, to red = 1.5.

### Sub-clustering reveals the presence of 5 tenocyte subclusters

Re-clustering of tenocyte populations alone identified 5 subclusters (Figure 3A), 3 of which localised to the IFM, “IFM_A”, “IFM_B”, and “IFM_C”, and 2 that were found in the fascicular matrix, “FM_A” and “FM_B”, based on the initial IFM or FM identity of re-clustered cells (Figure 3B) and on marker similarity with either the IFM or FM cluster, including DE of ECM genes *COMP, LOX, PRG4, TNXB, COL14A1, FAP* (Figure S4). Subclusters IFM_A and FM_A mainly comprised of cells from the original IFM cluster and FM cluster, respectively, whilst subclusters IFM_B, IFM_C, and FM_B mainly comprised of cells originating from the MixT cluster (Figure 3B). The top markers of each subcluster (most highly expressed markers) revealed markers for the FM_A subcluster were mainly associated with ECM production whereas FM_B top markers showed a high number of mitochondria-related genes. Further analysis of cluster FM_B, omitting the mitochondria-related genes in case they were masking other less expressed genes with significant functions, revealed markers of FM_B cluster were associated with response to stress. Markers for the IFM_A cluster showed gene expression related to the ECM, signalling in inflammation, and ECM remodelling. IFM_B top markers were associated with extracellular exosomes and IFM_C showed higher expression of genes related to immune function (Figure 3C).

**Figure 3.**
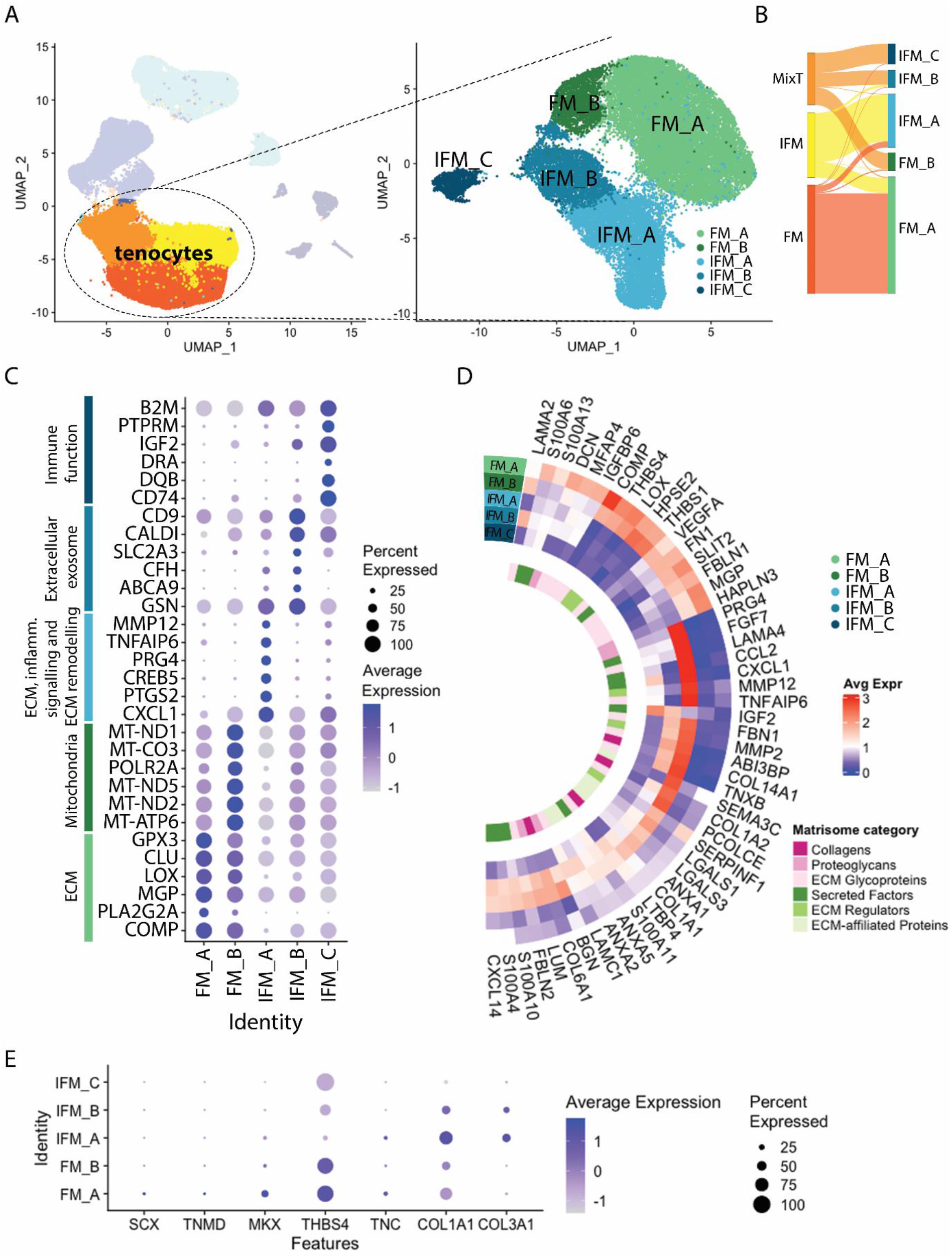
Sub-clustering reveals the presence of 5 tenocyte subclusters. (A) Tenocyte subclusters were identified as FM_A, FM_B, IFM_A, IFM_B, and IFM_C, based on their marker expression. (B) Sankey diagram showing the provenance of each subcluster cell in relation to the original clusters. (C) Dot plot showing the top 5 differentially expressed markers and associated functions in each subcluster. Scale indicates average expression and ranges from grey = -1 to blue = >1, dot size indicates the percentage of cells expressing the gene. (D) Heatmap showing average expression of the top 50 DE matrisomal genes across tenocyte subclusters in each tenocyte subcluster, with matrisome category indicated (circlize package, (Gu et al., 2014)). Scale indicates average expression and ranges from blue = 0, to white = 1, to red = 3. (E) Dot plot of established tenocyte lineage genes and their expression across the tenocytes subclusters. *MKX* and *THBS4* are predominantly expressed in FM subclusters whilst *COL1A1* and *COL3A1* in IFM subclusters. Scale indicates average expression and ranges from grey = -1 to blue = >1, dot size indicates the percentage of cells expressing the gene.

Analysis of matrisomal genes in young tenocyte clusters revealed distinct matrisome expression patterns between FM and IFM clusters, with clusters FM_A and IFM_A having the highest matrisome gene expression and IFM_C having the lowest matrisome gene expression (Figure 3D). FM clusters had the highest expression of *DCN*, glycoproteins *COMP, THBS1*, and *THBS4*, and ECM-related genes *LOX* (cross-linking) and *HPSE2* (matrix remodelling). IFM clusters had highest expression of collagens such as *COL1A2, COL14A1* (exclusive to the IFM) and *COL6A1* as well as proteoglycans *BGN* and *PRG4* and glycoproteins *TNXB* and *FBN1*. IFM clusters also showed higher gene expression of calcium binding proteins *S100A11, S100A4*, and *S100A10* which are involved in cell cycle progression and differentiation as well as cell motility and, when secreted, stimulation of cytokine production (Gross et al., 2014, Cerezo et al., 2014). Cluster IFM_C had the lowest matrisome expression but highly expressed secreted factors *IGF2* and *CXCL1*. Interestingly, FM and IFM subclusters had differential expression of laminin subunits with *LAMA2* expressed by FM subclusters (mostly FM_B) and IFM_B whilst *LAMA4* and *LAMC1* were expressed by IFM subclusters (mainly IFM_A and IFM_B, respectively). Similarly, fibulin expression revealed compartment-specific expression with *FBLN1* mostly expressed in FM clusters and *FBLN2* in IFM subclusters.

Expression of classical tenocyte lineage genes revealed *MKX* and *THBS4* expression in both FM subclusters and *COL1A1* and *COL3A1* expression in IFM_A and IFM_B clusters. *SCX* and *TNMD* only showed trace expression in FM_A subcluster (Figure 3E). The more recently identified tenocyte marker IGFBP6 (Turlo et al., 2019) also showed predominant expression in the FM_A subcluster, with lower expression in subclusters IFM_A and IFM_B.

### Tenocyte ageing is predominantly observed in IFM subclusters

There were no significant changes in the proportional size of tenocyte subclusters with ageing despite the apparent decrease in IFM_C subcluster size in old samples and concurrent increase in IFM_B subcluster size, as a percentage of total cells (Figure 4A). IFM_A subcluster appeared to change shape without any significant size change. An elongated tip at the bottom of the IFM_A cluster (Figure 4B) with presence of less differentiated cells (predicted, CytoTRACE) (Figure 4E and F) and expression of *POSTN* and *TPPP3* (Figure 4G) which have been associated with a progenitor phenotype in tendon cells (Harvey et al., 2019, Wang et al., 2021) is observed in young samples whilst absent from old samples (Figure 4B,E-G). Concurrently, expression of *POSTN* and *TPPP3* identified/localised to the IFM_A cluster was significantly decreased with ageing (Figure 4G). IFM subclusters also showed the most DE genes with ageing, particularly subcluster IFM_A (Figure 4C), with DE genes related to the ECM, signalling in inflammation, and ECM remodelling. Subcluster IFM_A also had the most significantly DE core matrisome genes (Figure 4D). Differential expression of core matrisome genes further highlighted a decrease in collagen gene expression in aged tenocytes (*COL1A1, COL3A1, COL6A1-3, COL14A1*) with the exception of *COL4A1-2*, which increased in the IFM subclusters specifically. Other genes encoding important components of the tendon ECM, such as proteoglycans *ASPN, BGN* and *PRG4* and glycoprotein *TNXB*, were generally downregulated in aged tenocytes. *FBN1* and *MFAP5*, which are associated with the elastic properties of tendons, were also downregulated in all tenocyte clusters with ageing whilst *SPP1* (encoding osteopontin) which is involved in tendon matrix remodelling (Mori et al., 2007) was upregulated. Interestingly, genes encoding proteins with an important role in fascicular matrix composition such as glycoproteins *COMP* and *THBS4*, proteoglycan *DCN* and cross-linking *LOX*, appeared to decrease in the FM subclusters and increase in the IFM subclusters (Figure 4D and Figure S5). Finally, *LAMA2* expression increased in all tenocyte subclusters whilst *LAMA4* decreased in all subclusters.

**Figure 4.**
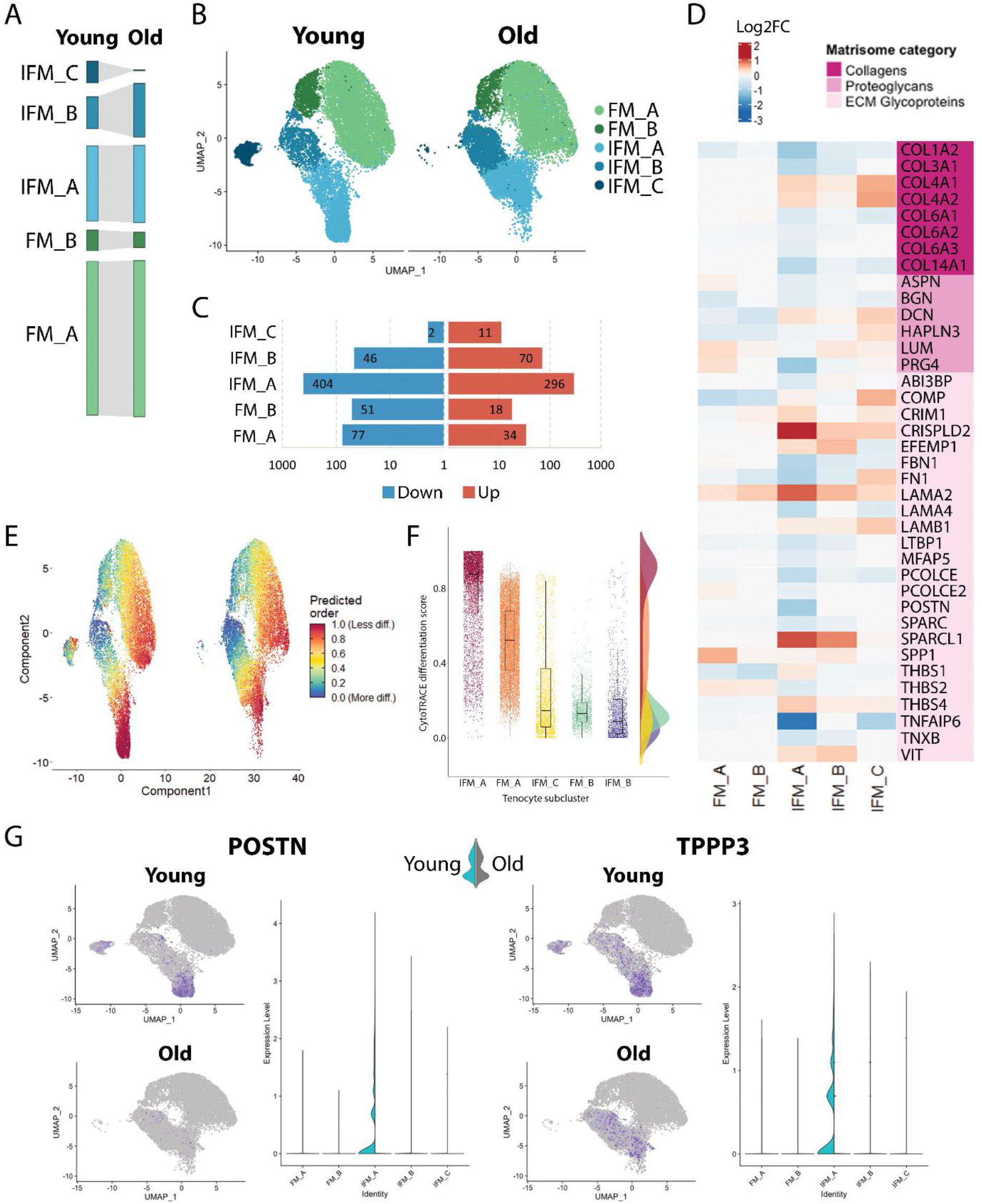
Tenocyte ageing is predominantly observed in IFM subclusters. (A) The percentage of cells in the subclusters was not statistically significantly affected by ageing, despite an apparent decrease in IFM_C cell number. (B) UMAP of tenocyte subclusters in young and old tendons; the distribution of cells in subcluster IFM_A changes with ageing. (C) The number of DEGs with ageing varied between subclusters, with the highest number of DEGs in the IFM_A clusters. Data are plotted on a log_10_ scale. (D) Heatmap showing DE of selected core matrisome genes with ageing in each tenocyte subcluster. Scale indicates log_2_FC and ranges from blue = –3, to white = 0, to red = 2. (E) UMAP of tenocyte subclusters with prediction of differentiation state in the young and old tenocytes and (F) raincloud plot of tenocyte subclusters in young samples by order of least differentiation (CytoTRACE). Subcluster IFM_A has the largest number of least differentiated cells in young samples (raincloud plot and UMAP), which are located particularly in the elongated tip at the bottom of the subcluster. Whereas in old samples, this elongated tip of the IFM_A subcluster is absent and a reduction in the number of least differentiated cells is noted. Scale ranges from blue = more differentiated to red = least differentiated through green, yellow and orange. (G) UMAP and violin split plot of *POSTN* and *TPPP3* expression in tenocyte subclusters in young and old tendons. *POSTN* and *TPPP3* expression, which has been associated with a progenitor phenotype in tendon cells, is observed in young samples in the elongated tip at the bottom of subcluster IFM_A and it is significantly decreased with ageing. Scale indicates expression and ranges from grey = 0 to blue = 3.

### FM and IFM tenocyte clusters become the primary sources of outgoing signalling in ageing

To examine the cell communication between different cell types, we performed cell-to-cell communication analysis using “CellChat”, a package that uses manually curated literature-supported ligand-receptor interactions (Jin et al., 2021). Here, we focused on “secreted signalling” interactions in order to avoid overcrowding due to the abundance of ECM-cell interactions and cell-cell interactions. We unveiled 230 secreted signalling interactions in young tendon cells and 238 secreted signalling interactions in old tendon cells (interaction strength 1.939 and 2.326 respectively). Mural cells (MuC1), followed by the tenocytes (FM, IFM), predominated in outgoing signalling in young tendon, whilst incoming signals were mainly received by vascular endothelial cells (EC-V) followed by macrophages (Mφ) and T cells (TC) both in young and old tendon (Figure 5A). With ageing, the FM and IFM tenocyte clusters showed the largest change in outgoing secreted signalling, increasing their signalling interaction strength and becoming the primary sources of outgoing signalling in old tendon cells. More specifically, the FM and IFM tenocytes clusters increased their signaling interaction strength to all clusters apart from the mixed tenocyte cluster (MixT), with signals to the mural cells (MuC2) and T cells (TC) showing the biggest increase. These clusters, MuC2, TC and MixT, were also the target clusters that showed the most changes in incoming secreted signalling with ageing, with the MuC2 and TC clusters showing increased incoming signalling interaction strength with ageing and the MixT cluster decreased interaction strength (Figure 5A).

**Figure 5.**
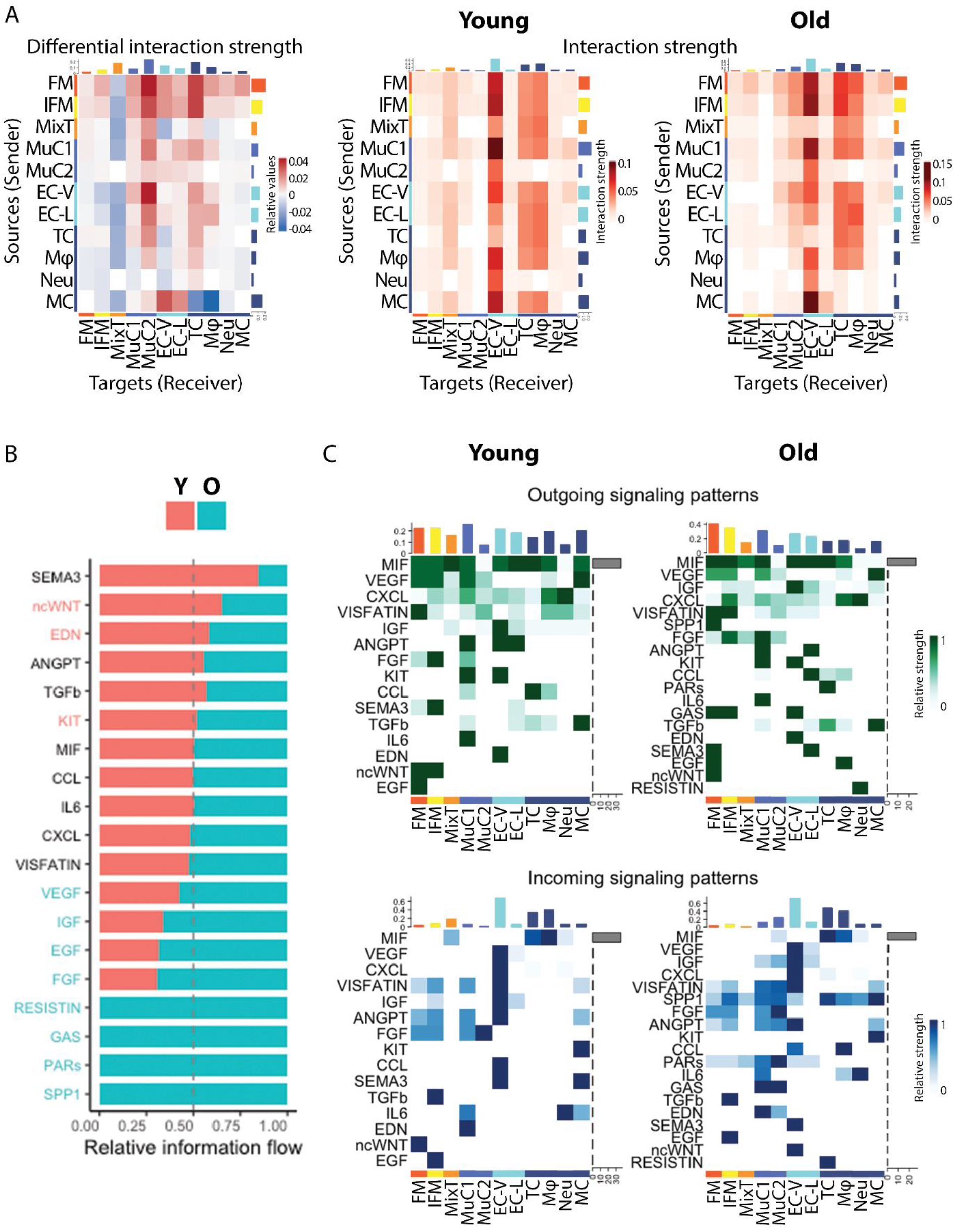
FM and IFM tenocyte clusters become the primary sources of outgoing signalling in ageing. (A) Heatmap of differential secreted signalling interaction strength among tendon clusters following ageing, along with the heatmaps of secreted signalling interaction strength among clusters in the young and old tendons. With ageing, the FM and IFM tenocyte clusters showed the largest change in outgoing secreted signalling, increasing their signalling interaction strength and becoming the primary sources of outgoing signalling in old tendon cells. Scales indicate relative values and interaction strength and range from blue to red and from white to red, respectively. (B) Secreted signalling pathways enriched in young (red) and old (blue) tendon. (C) Outgoing and incoming signalling patterns for each secreted signalling pathway and cluster in young and old tendon. Scales indicate relative strength and range from white to green and from white to blue.

Pathways such as non-canonical WNT were downregulated with ageing whilst others such as the growth factor pathways VEGF, IGF, EGF, FGF were upregulated in old tendon (Figure 5B). In particular, FM tenocytes in young tendons exerted a strong signal in Visfatin, ncWNT, and EGF signalling, IFM tenocytes in FGF, SEMA3, and ncWNT signalling, and MixT cells in MIF signalling (Figure 5C). On the other hand, FM tenocytes in young tendons received a strong signal from ncWNT signalling and IFM tenocytes from TGFb and EGF signalling. Tenocyte-related enriched secreted signalling pathways that altered with ageing were the ncWNT pathway, which was enriched in young cells and the growth factor pathways VEGF, IGF, EGF, FGF, and the GAS, PARs, and SPP1 pathways, which were enriched in old cells. ncWNT signalling in young tendon was exerted from FM and IFM tenocytes to FM tenocytes through *WNT11*, and EGF signalling, which was enriched in old tenocytes, was from FM to IFM tenocytes and was through *HBEGF*.

## Discussion

This study has unveiled the complexity of cell populations within the tendon IFM for the first time and identified specific populations that are disproportionately affected by ageing. The age-related dysregulation of IFM tenocytes likely has important implications for maintenance of tendon homeostasis in aged individuals and may contribute to the impaired reparative capacity and subsequent increased risk of tendon injury with ageing.

Our results demonstrate that the IFM houses a unique tenocyte population that can be distinguished from fascicular tenocytes due to higher expression of *PRG4* and *TNXB*, and lower expression of *COMP, LOX* and *THBS4*. Tendon endothelial, mural and immune cell populations also localise to the IFM, supporting recent work that has identified the presence of an endothelial-like basement membrane in the IFM (Marr et al., 2022). Previous scRNAseq studies of mouse and human tendons have identified a range of similar cell populations to those identified in the current study, encompassing tenocytes, endothelial cells and immune cells (Kendal et al., 2020, De Micheli et al., 2020). While these studies have unveiled the heterogeneity of tendon cell populations, the majority of murine tendons lack an identifiable IFM. Furthermore, ageing processes differ between long-lived and short-lived animals, with continual growth of mice throughout their lifespan, making the mouse an unsuitable model to study the IFM in ageing tendon (Jilka, 2013, Lee and Elliott, 2019). It is very difficult to collect viable, healthy tendon tissues from young human donors making studies on human tendons impractical. Due to the similarities in tendon pathophysiology between humans and horses and tendon injuries occurring preferentially in elastic tendons with a prominent IFM such as the equine SDFT, the SDFT is therefore a highly appropriate model in which to study the effect of ageing on IFM cell populations. Limitations associated with this model are that the sex, breed and exercise history of the horses are unknown, which may account for the variability between individuals seen in the data, as well as the incomplete annotation of the equine genome. Despite these limitations, we were still able to identify a host of age-related alterations in gene expression between different cell clusters.

We identified three tenocyte clusters, two of which localised to the IFM and FM respectively. The third tenocyte cluster was classed as a mixed tenocyte population (MixT) due to the presence of cells with DE of genes associated with both IFM and FM tenocytes. This clustering was driven by downregulation of ribosomal genes in the mixed tenocytes, which suggests a decreased synthetic activity in these cells (Steffen and Dillin, 2016). Further analysis of tenocyte subclusters revealed distinct expression of matrisomal genes, particularly between IFM and FM subclusters, reflecting the differences seen in matrix composition between tendon IFM and FM regions. The low expression of established tenocyte markers *SCX, MKX* and *TNMD* in these subclusters is supported by single-cell sequencing data from human and mouse tendons, which also demonstrate limited expression of these markers in adult tenocytes, suggesting that *SCX, MKX* and *TNMD* expressing cells have a more important role in tendon development than in mature tissue (Kendal et al., 2020, De Micheli et al., 2020). Average *COL1A1* expression was higher in IFM subclusters compared to FM subclusters, which may seem counterintuitive due to the enrichment of type I collagen in the fascicular matrix (Thorpe et al., 2016b). However, our previous studies have shown that collagen is turned over more rapidly within the IFM, likely accounting for the increased *COL1A1* gene expression measured within IFM tenocytes (Choi et al., 2020). One FM subcluster showed higher expression of mitochondrial-related genes compared to the other tenocyte subclusters. High mitochondrial gene expression has been previously reported in a subset of tenocytes in response to mechanical stimulation and in association with cellular stress (Still et al., 2021) and during early healing of tendon in relation to hypoxia in tenocytes (Thankam et al., 2018), but also in dying cells or cells undergoing apoptosis (Still et al., 2021). Here, the FM subcluster showing high mitochondrial gene expression also showed differential expression of genes related to cellular stress but not cell death or apoptosis. As we filtered out dead and dying cells prior to single cell sequencing, we are confident that the high expression of mitochondrial genes in this cluster is not due to cell death induced by sample processing.

The putative tendon progenitor markers *TPPP3* (Harvey et al., 2019) and *POSTN* (Noack et al., 2014, Wang et al., 2021) were also predominantly expressed in the largest IFM subcluster, in particular in a region of the cluster where less differentiated cells were observed, corroborating a less differentiated state for these cells. With ageing, *TPPP3* and *POSTN* expression was significantly decreased in IFM tenocytes, as were the number of IFM tenocytes in a less differentiated state, which may provide some insight into the reduced ability for regeneration in aged tendons. Decreased levels of both *TPPP3* and *POSTN* have been reported in other cell types with ageing and senescence (Egbert et al., 2014, Schwartz et al., 2021) supporting a putative role for these genes in cell and tissue ageing. *TPPP3* expression, however, has also been reported in a tendon immune cell population in mouse tendon (De Micheli et al., 2020), indicating these cells may not be true progenitors and as such these markers warrant further investigation in tendon and across species.

One of the key findings of the current study is the age-related alteration of IFM cell populations, with loss of cluster-differentiating gene expression signatures in IFM and MixT tenocyte and mural cell clusters. Similar results have been found in ageing skin fibroblasts, with loss of cellular identity and functional priming and reduced cell interactions with increasing age (Solé-Boldo et al., 2020). Ageing IFM cell populations also showed dysregulation of senescence-associated genes. Previous studies have demonstrated that human and rodent Achilles tendon-derived stem/progenitor cells express several markers of senescence with increasing age (Kohler et al., 2013, Chen et al., 2020). However, no studies have investigated senescence in other tendon cell populations, and this remains an important area for future research. We also identified alterations in a panel of inflammation-associated genes with ageing, particularly in IFM tenocytes and mural cells, indicating a deregulated ability to modulate inflammation, which may result in the inflamm-aging phenotype that is seen in aged tendons during injury (Dakin et al., 2012). A panel of genes associated with loss of proteostasis were also DE with ageing. This supports our previous findings, which have demonstrated decreased turnover within the IFM specifically (Thorpe et al., 2016b). IFM tenocytes were also the only cluster to show age-related changes in cell cycling, with an increase in the percentage of cells in S phase in aged tendons without a concomitant increase in the G2/M phase. Alterations in the duration of the S phase have been previously noted in senescence and following replicative stress (Prieur et al., 2011, Ercilla et al., 2016) but further investigation is required to elucidate the events leading to the observed increase of cells in S phase and its implications for aged tendon.

Results showed alterations in expression of matrisomal genes with ageing in tenocytes, with a general decrease in expression of collagens, with the exception of *COL4A1&2*, which increased in IFM subclusters. These genes encode an integral component of the IFM basement membrane (Marr et al., 2022), and therefore these changes may result in alterations to IFM basement membrane structure. In addition, the decrease in *LAMA4* expression with ageing in tenocytes may have important implications for healing, as it has previously been shown that LAMA4 may be important for the recruitment of IFM cell populations after injury (Marr et al., 2021). A decline in the quality of the tendon ECM with ageing was also evidenced across IFM and FM regions. Decrease in expression of matrisomal genes, such as *BGN, ASPN, PRG4, FBN1, MFAP5* associated with IFM composition and mechanical properties, was noted in IFM tenocytes, whilst *COMP* and *LOX*, associated with FM composition and properties decreased specifically in FM tenocytes (Thorpe et al., 2016b, Godinho et al., 2017, Zamboulis et al., 2020). The decline in ECM integrity is considered not only to be fundamental to the functional impairment in tendon but is also a driver of cellular ageing and disease progression (Ewald, 2020) and as such, of critical importance in ageing tendon.

Mural cell ageing may also have important implications for tendon function. Few studies have investigated the effect of microvascular cell ageing on tendon function, however studies have shown that peritendinous blood flow in the Achilles tendon is lower in aged individuals compared to young individuals (Langberg et al., 2001). There is also a decrease in the number of capillaries within healthy tendon with ageing, and neo-vascularisation is reduced in old rats following injury (Kannus et al., 2005, Márquez-Arabia et al., 2017, Riggin et al., 2022). These changes may be driven by altered mural cell function due to the dysregulated gene expression described in the current study as well as altered cellular environment. Indeed, age-related microvascular dysfunction is common in other tissues including skeletal muscle (Scioli et al., 2014), with senescence being preceded by vascular attrition in a range of tissues (Chen et al., 2021). Pericyte functionality is also diminished with ageing, with impaired regenerative capacity in skin (Zhuang et al., 2021).

Analysis of cell-to-cell communication revealed a complex signalling network between tendon cell populations that was altered with ageing, particularly in tenocytes, mural cells, and T cells. This alone highlights the importance of elucidating the heterogeneity of tendon cell populations and the importance of using co-culture systems to recapitulate complex in vivo interactions. Of note, SPP1 and EGF signalling through *HBEGF*, which have been reported to participate in the active remodelling of tendon during injury repair (Ackerman et al., 2021), were enriched in aged tendons.

The discovery of the extensive heterogeneity of tendon cell populations has important implications for studying tendon-derived cells *in vitro*. The response of tendon-derived cells to a wide variety of physicochemical stimuli has been extensively characterised *in vitro* (Wall et al., 2016, Ryan et al., 2021, Wang et al., 2018), however, these studies have typically assumed that tendon-derived cells are a homogenous population of fibroblastic-like tenocytes, utilising culture conditions and experimental conditions appropriate for such a population, potentially impacting the relevance of such results to *in vivo* tendon cell behaviour. Identification of markers for each tendon cell population will allow future studies to use cell sorting approaches to study responses of individual populations in isolation or in specific combinations, which will provide much more in-depth information on the role of each population in tendon homeostasis, ageing and injury. However, it will be important to optimise culture conditions to ensure phenotypic stability is maintained in these different cell populations.

While this study has identified novel IFM-localised cell populations that are prone to age-related dysfunction, there is a pressing need to define the mechanisms that are driving these age-related alterations. Previous studies have demonstrated that the IFM is disproportionately affecting by ageing, with stiffening occurring, likely due to accumulation of microdamage and reduced protein turnover (Thorpe et al., 2013, Godinho et al., 2017, Thorpe et al., 2016b). These changes will likely alter the cell microenvironment, which could negatively affect cell phenotype and lead to the age-related changes observed in the current study. Additionally, the accumulation of non-enzymatic modifications within extracellular matrix proteins, which are common in ageing tendon, have been found to induce cell senescence and tissue fibrosis (Fedintsev and Moskalev, 2020, Thorpe et al., 2010, Selman and Pardo, 2021). Alternatively, there may be intrinsic changes within IFM cell populations driving age-related alterations in cell phenotype. This remains an important area for future research to allow development of therapeutics that can effectively limit and/or reverse tendon age-related dysfunction and subsequent injury.

## Conclusions

We have uncovered the heterogeneity of IFM-localised cell populations within tendon, revealing the presence of diverse cell types including tenocytes, microvascular cells and immune cells. The IFM cell populations are disproportionately affected by ageing, with dysregulation of genes associated with senescence, proteostasis and inflammation, making these cells likely targets when developing more effective therapeutics for age-related tendon injury.

## Materials and Methods

### Sample acquisition and preparation

Forelimbs, distal to the carpus, were collected from young (n=4; age range:3-4 years) and aged (n=4; age: >17 years) horses euthanised for reasons unrelated to this project from a commercial abattoir. Sample collection was approved by the Royal Veterinary College’s Clinical Research Ethical Review Board (URN 2016 1627B). Two ∼3 cm pieces of superficial digital flexor tendon (SDFT) were harvested from the mid-metacarpal region of one forelimb from each horse on the same day as euthanasia. One piece was snap frozen in isopentane in liquid nitrogen for subsequent histology and spatial analysis of cell niches, and the other was digested to obtain cell populations as described below.

### Tendon digestion and generation of single cell suspensions

SDFT pieces were washed several times in Dulbecco’s Modified Eagle’s Medium (DMEM) with antibiotics and antimycotics, the epitenon was removed and the tendon core was finely minced (a few mm^3^) under sterile conditions. Samples were digested (3 mg/mL collagenase type 2, 1 mg/mL dispase, 100 μg/mL DNase I in DMEM supplemented with 50 U/mL penicillin, 50 μg/mL streptomycin, 0.5 μg/mL amphotericin B and 10 % foetal bovine serum) for 4 h at 37° C with agitation. Samples were strained (40 μm filters) and dead cells removed using a dead cell removal kit according to the manufacturer’s instructions (Miltenyi Biotec). Cell viability was determined using trypan blue staining and cells were counted using a haemocytometer, followed by resuspension in 0.4 % BSA in PBS at ∼1000 cells/μl. Viability was ≥ 90 % for all samples.

### Single cell RNA sequencing

Approximately 6,500 cells from each sample were prepared for scRNAseq using a Chromium single cell 3’ reagent kit (10x Genomics) according to the manufacturer’s instructions. Libraries were pooled and sequenced on an Illumina® NovaSeq 6000 (Illumina®, San Diego, USA) on an SP flow cell, generating 28 bp x 91 bp paired-end reads. Data have been deposited at EMBL-EBI under the project ID PRJEB57256.

### Bioinformatic Analysis

Reads were aligned using Cell Ranger (v.6.0.0) on horse genome (EquCab3.0.103). Barcode swapping events and empty droplets were excluded using DropletUtils (Griffiths et al., 2018) and subsequent downstream analysis was performed using Seurat (v4.1.1) in R Studio (v2021.09.2) (Butler et al., 2018). Quality control was performed as follows: filters: >500 unique molecular identifiers/cell; 250-5500 genes/cell; Log10Gene Per UMI >0.8; <10% mitochondrial reads/cell. Any genes expressed in less than 10 cells were excluded from subsequent analysis. Following data normalization by SCTransform (Hafemeister and Satija, 2019), principal component analysis and integration was performed using Harmony (Korsunsky et al., 2019). The ‘FindCluster’ function was used to identify cell clusters (0.2 resolution). Post-clustering, further manual quality control revealed the presence of one cluster comprised of cells originating predominantly from one sample. Further analysis of this cluster indicated it likely consisted of cells derived from the epitenon and therefore it was excluded from any further analysis. Genes differentially expressed (DE) between remaining clusters were identified using the ‘FindAllMarkers’ function in Seurat.

### Analysis of age-related alterations

Genes differentially expressed (DE) between age groups within each cluster were identified using the ‘FindAllMarkers’ function. Of these DE genes, those associated with ageing were identified and classified using the Ageing Atlas (Consortium, 2020). The top 25 genes in each cluster were compared between young and old to identify the number of genes conserved with ageing. To assess changes in inflammatory gene signatures, DE genes with ageing associated with the GOTerm ‘inflammatory response’ (GO:0006954) (Smith and Eppig, 2009) were identified.

### Analysis of tenocyte clusters

The tenocyte clusters IFM, FM, and MixT, were re-clustered (0.1 resolution) and genes DE between subclusters and with ageing identified as described above. The top 50 DE matrisome-associated genes between subclusters were identified and classified according to matrisome categories (Shao et al., 2019), and DE matrisome-associated genes with ageing were identified. Heatmaps with circular layout were created using the R package “circlize” (Gu et al., 2014).

### Cell differentiation state analysis

Cell differentiation state analysis was carried out with the open source package CytoTRACE (Cellular (Cyto) Trajectory Reconstruction Analysis using gene Counts and Expression) (Gulati et al., 2020); a computational method that predicts the differentiation state of cells from single-cell RNA-sequencing data based on the number of detectably expressed genes per cell, or gene counts which are determinants of developmental potential. The differentiation state of tenocytes was visualised on UMAPs of young and old re-clustered tenocytes and raincloud plots (Allen et al., 2021), ranking the order of differentiation state for young re-clustered tenocytes.

### Cell communication analysis

Cell-to-cell communication analysis was performed using the open source R package CellChat (Jin et al., 2021) focusing on secreted signalling interactions also termed paracrine/autocrine signalling interactions. To infer the cell cluster-specific communications, CellChat identified over-expressed ligand-receptor interactions by identifying over-expressed ligands or receptors in cell clusters. CellChat was then used to compare cell-cell communication between young and old samples to identify interactions between cell clusters that were significantly changed with ageing, alongside identifying how signalling sources and targets changed. In addition, CellChat identified significantly decreased or increased signalling pathways in ageing by comparing the information flow for each signalling pathway; the information flow is defined by the sum of communication probability among all pairs of cell groups in the inferred network. Finally, outgoing and incoming signalling pathways were visualised for young and old samples to identify age-related differences in pathways for each cell cluster. Heatmaps were used to illustrate the cluster communications and pathways along with their differences with ageing.

### Spatial distribution

Longitudinal cryosections were cut from the snap-frozen SDFT samples at a thickness of 20 μm and adhered to glass slides. Haematoxylin and eosin staining was performed using standard protocols to ensure that all samples displayed a normal morphology and were free from any microscopic signs of injury. Immunohistochemistry was performed as described previously to localise markers for each cell cluster; details of primary and secondary antibodies, and blocking conditions are shown in Table 1 (Marr et al., 2021). Briefly, tendon sections were thawed, fixed in ice cold acetone:methanol (50:50) for 5 minutes, washed with tris-buffered saline (TBS), blocked and then incubated in primary antibody overnight at 4° C. Following TBS washes and hydrogen peroxide treatment (0.3 %, 15 min), secondary antibodies were applied at room temperature for 1 hour. Staining was developed with 3,3’-diaminobenzidine (5 min, Vector Labs, San Francisco, CA, USA), and sections were counterstained with haematoxylin for 30 s, dehydrated (Gemini AS Automated Slide Stainer, Thermo Scientific) and mounted with DPX. Sections were cured overnight and imaged using brightfield microscopy (DM4000B upright microscope; objectives: 10× HC PL FLUOTAR PH1, 20× HC PL FLUOTAR PH2; DFC550 colour camera; LAS-X version 3.7 software (Leica Microsystems)).

**Table 1.** Details of primary and secondary antibodies, and blocking conditions used for immunolabelling experiments.

| Primary Antibody | Supplier | Dilution | Secondary Antibody | Supplier | Dilution | Blocking conditions |
| --- | --- | --- | --- | --- | --- | --- |
| LOX | Novus Biologicals (NB100-2530) | 1:50 | Goat anti-rabbit IgG | Dako | 1:500 | 5% goat serum; 5% horse serum; 1% BSA |
| MYH11 | Thermo Scientific (PA5-82526) | 1:200 | Goat anti-rabbit IgG | Dako | 1:500 | 5% goat serum; 5% horse serum; 1% BSA |
| PECAM1 | Abcam (ab28364) | 1:50 | Goat anti-rabbit IgG | Dako | 1:500 | 5% goat serum; 5% horse serum; 1% BSA |
| TNXB | St John's Laboratory (STJ95967) | 1:300 | Goat anti-rabbit IgG | Dako | 1:500 | 5% goat serum; 5% horse serum; 1% BSA |
| RGS5 | St John's Laboratory (STJ95440) | 1:600 | Goat anti-rabbit IgG | Dako | 1:500 | 5% goat serum; 5% horse serum; 1% BSA |
| CD74 | St John's Laboratory (STJ96829) | 1:100 | Goat anti-rabbit IgG | Dako | 1:500 | 5% goat serum; 5% horse serum; 1% BSA |

## Acknowledgements

The authors acknowledge the contribution of Shumeng Duan and Binyao Zhang, Physical Therapy in Musculoskeletal Healthcare and Rehabilitation MSc students at UCL, for their contributions to initial data analysis and the Centre for Genomic Research at the University of Liverpool for performing the sequencing.

## Author Contributions

DEZ – Conceptualization, Funding acquisition, Investigation, Methodology, Writing - original draft, Writing - review and editing, Visualisation, NM – Investigation, Visualisation, Writing - review and editing, LL - Investigation, Visualisation, Writing - review and editing, HLB – Conceptualization, Funding acquisition, Writing - review and editing, HRCS - Conceptualization, Funding acquisition, Writing - review and editing, PDC - Conceptualization, Funding acquisition, Writing - review and editing, CTT - Conceptualization, Funding acquisition, Investigation, Methodology, Writing - original draft, Project administration, Writing - review and editing, Supervision.

## Supplementary data

**Supplementary Figure 1.**
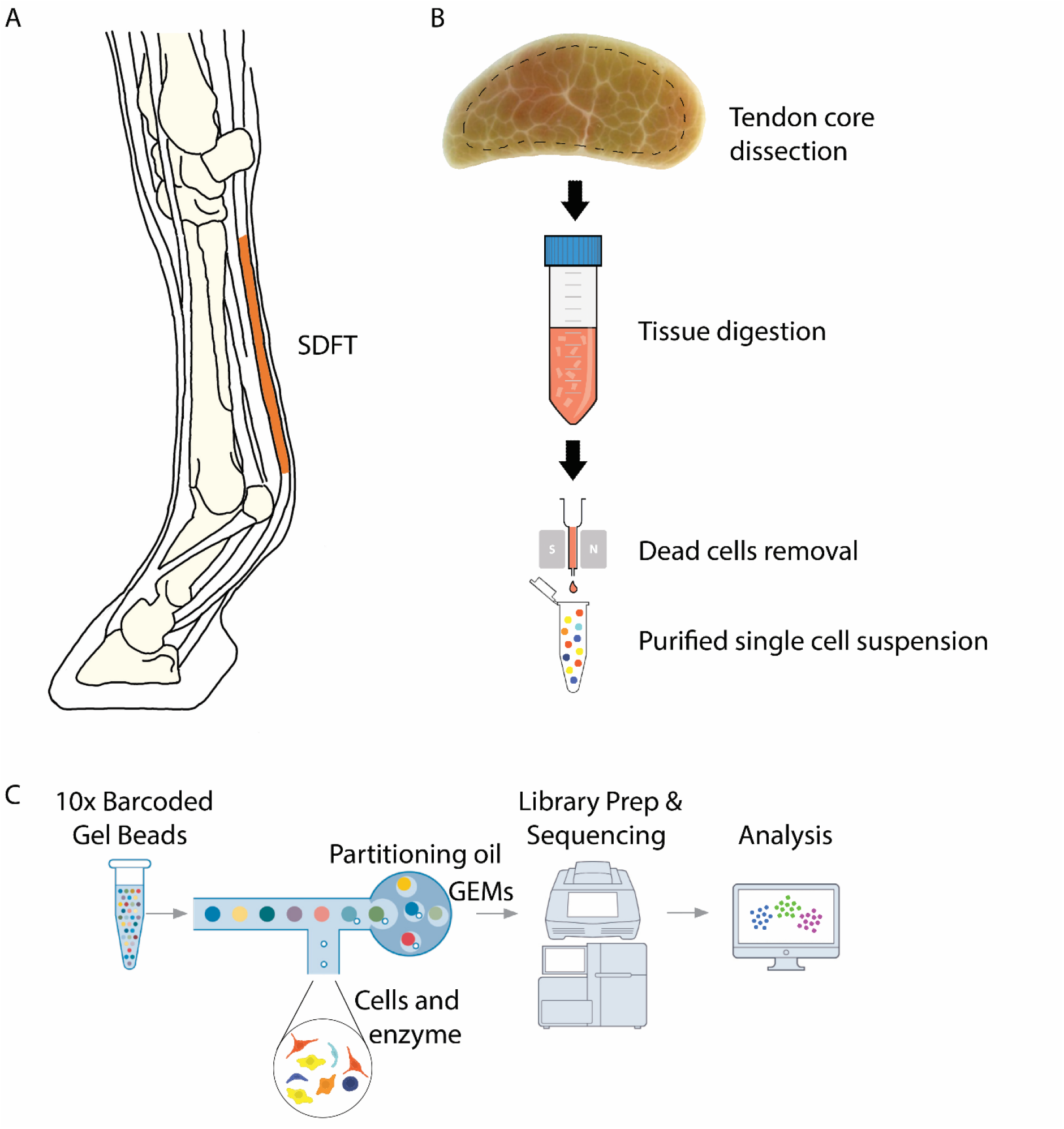
Schematic showing sample collection and processing for single-cell RNA sequencing. (A) Superficial digital flexor tendons (SDFT) were harvested from the forelimbs of young and old horses (n=4/age group). (B) Samples were collected from the core of the mid-metacarpal region of the tendon, finely diced and enzymatically digested. Dead cells were removed to generate single cell suspensions. (C) Single cells were encapsulated using gel beads in emulsion (GEM), followed by library preparation, RNA sequencing and data analysis.

**Supplementary Figure 2.**
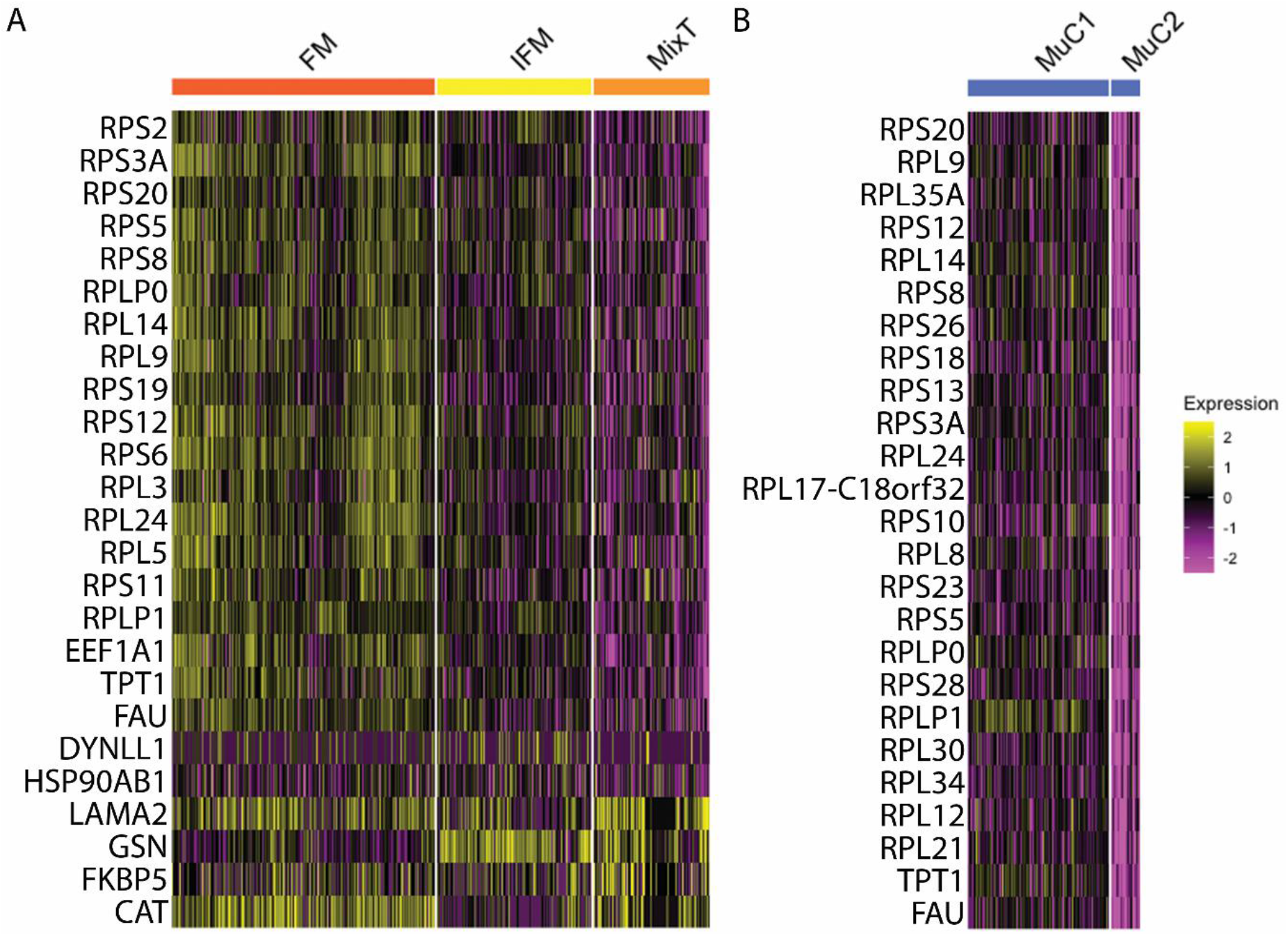
(A) Heatmap depicting gene expression of the top 25 markers for the MixT cluster (ROC analysis) across the FM, IFM and MixT clusters (the ROC analysis returns a ‘predictive power’ ranked matrix of putative differentially expressed genes based on the ability of these genes to classify cells in a cluster). The MixT cluster shows lower expression of ribosome biogenesis-specific genes compared to the FM and IFM clusters. (B) Heatmap depicting gene expression of the top 25 markers for the MuC2 cluster (ROC analysis) across the MuC1 and MuC2 clusters. The MuC2 cluster shows lower expression of ribosome biogenesis-specific genes compared to the MuC1 cluster. Scale indicates log_2_FC expression and ranges from pink = <-2 to yellow = >2.

**Supplementary Figure 3.**
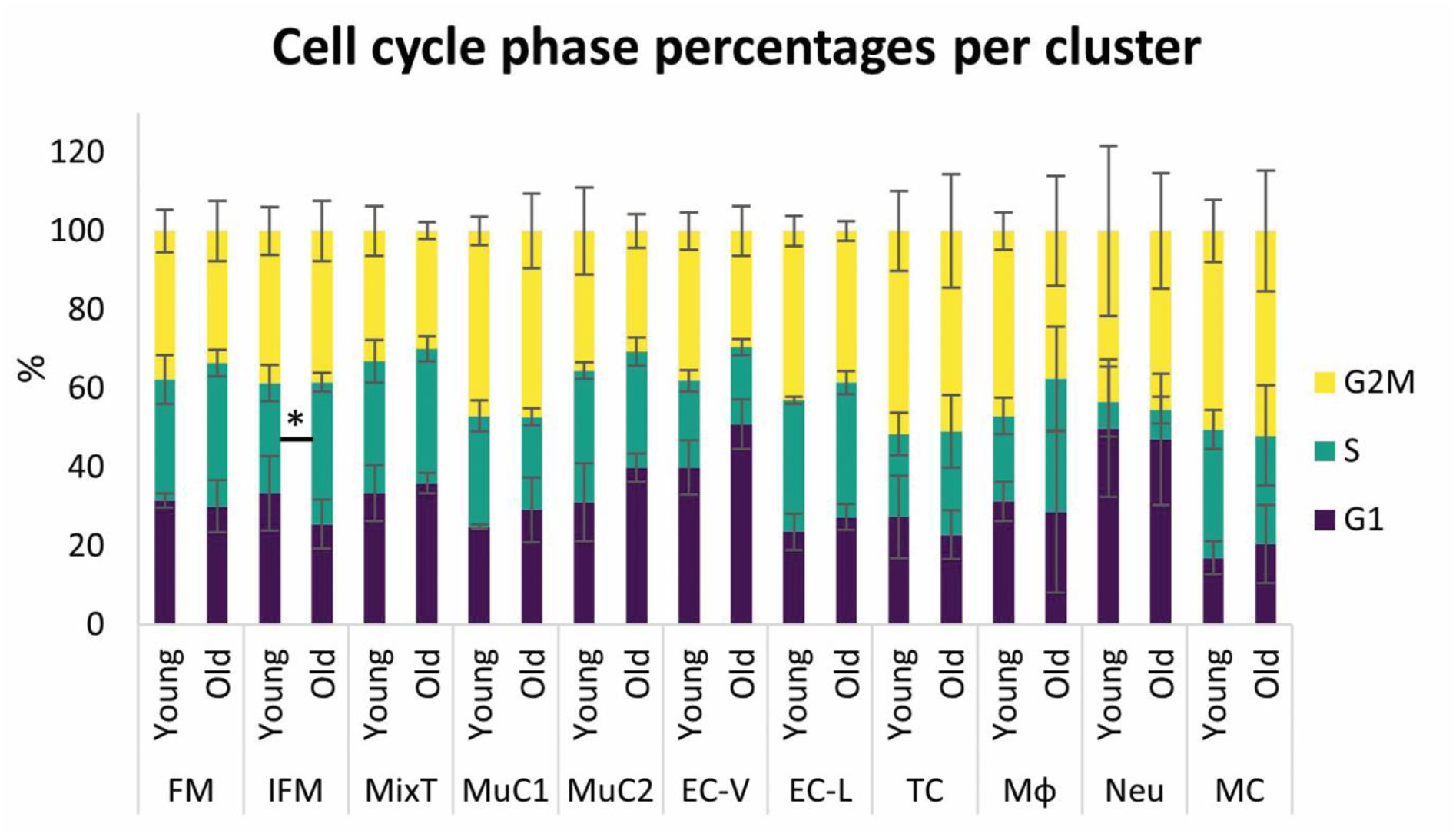
Percentage of cells classifying as G1, S, and G2M phase per cluster in young and old samples. The IFM tenocyte cluster was the only one to show an age-related effect on cell cycling with a significantly larger percentage of cells classified in the S phase in the old samples (p=0.023, t-test).

**Supplementary Figure 4.**
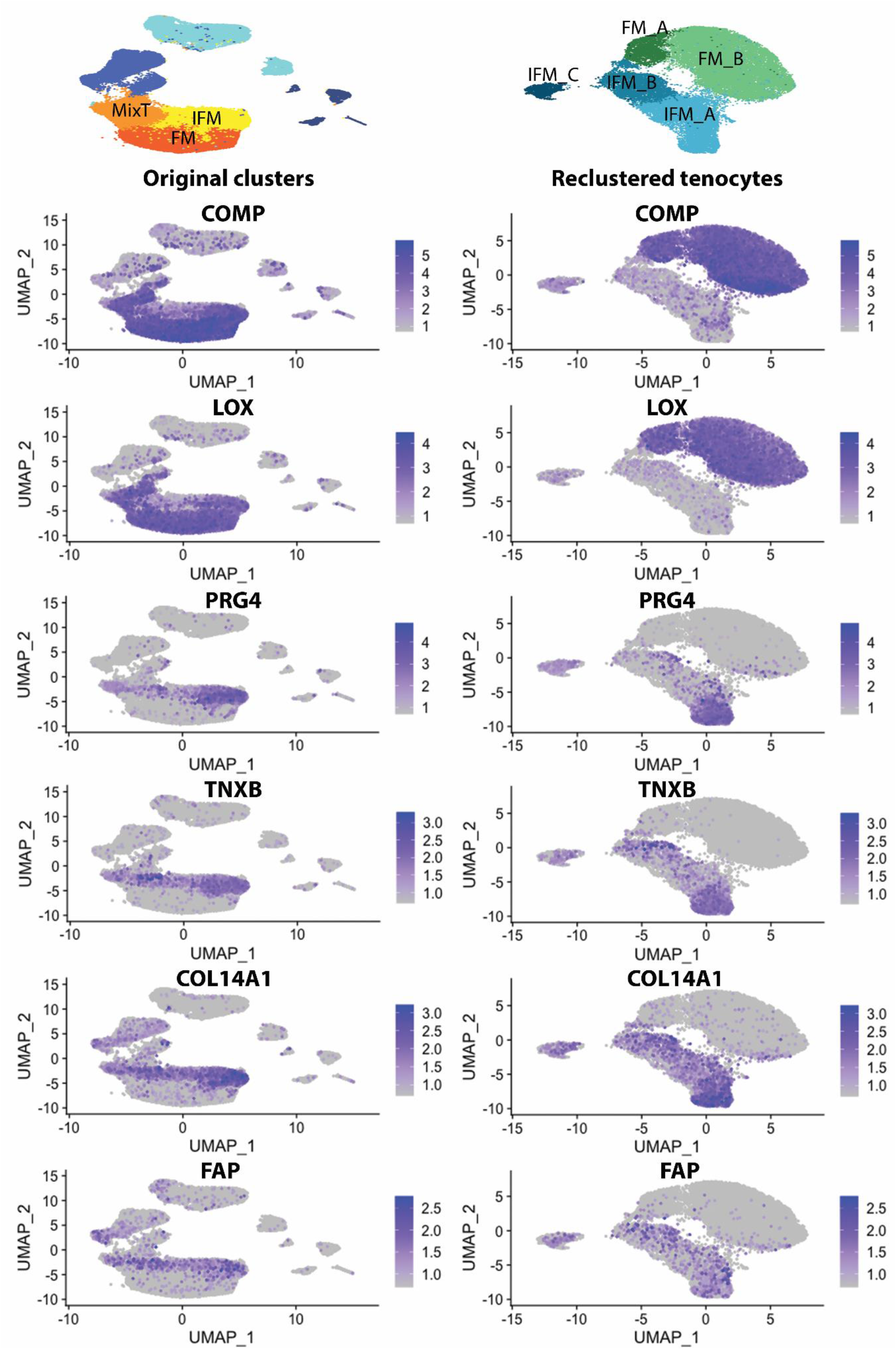
UMAPs of *COMP, LOX, PRG4, TNXB, COL14A1* and *FAP* expression in the original tenocyte clusters and the re-clustered tenocytes. Tenocytes subclusters “FM_A” and “FM_B” and “IFM_A”, “IFM_B”, and “IFM_C” were confirmed as FM and IFM tenocytes respectively based on their differential expression of FM tenocytes markers *COMP* and *LOX*, and IFM tenocytes markers *PRG4, TNXB, COL14A1*, and *FAP*. Scale indicates expression and ranges from grey to blue.

**Supplementary Figure 5.**
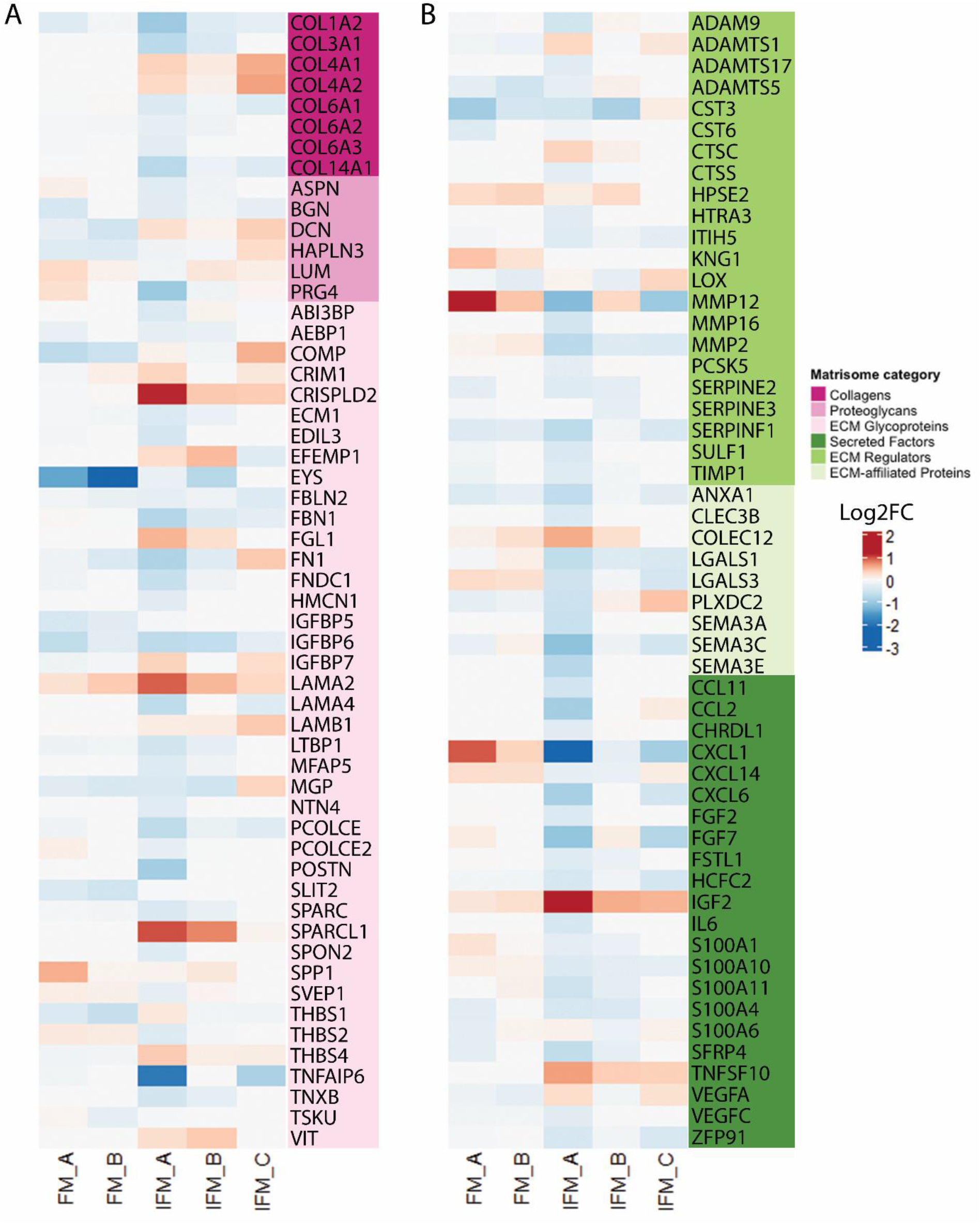
Heatmap of DE core matrisome (A) and matrisome-related (B) genes with ageing in each tenocyte subcluster. The core matrisome categories, “Collagens”, “Proteoglycans”, and “ECM Glycoproteins”, and matrisome-related categories, “Secreted Factors”, “ECM Regulators” and “ECM-affiliated Proteins”, are colour-coded. Scale indicates log_2_FC and ranges from blue = –3, to white = 0, to red = 2.

